# *Klebsiella pneumoniae* BolA contributes to cell morphology, siderophore production, stresses challenge, cell adhesion and virulence

**DOI:** 10.1101/2021.04.05.438546

**Authors:** Xiangjin Yan, Feiyang Zhang, Manlin Ding, Li Xiang, Jiawei Bai, Qin Li, Yingshun Zhou

## Abstract

*Klebsiella pneumoniae* infection is one of the important reasons for the increased of morbidity and mortality. The main virulence factors of *K. pneumoniae* include capsule polysaccharide, lipopolysaccharide, fimbriae, outer membrane proteins and siderophores. BolA homologues form a broadly conserved family of proteins in prokaryotes and eukaryotes. In *Escherichia coli, bolA* expression is quickly induced in response to different stresses or stationary phase that rapidly adapt to changing environments. In this report, we confirmed that *bolA* mutant strain exhibited increased sensitivity to bile and oxidative stresses. In addition, gene deletion showed that *bolA* has an important role for the adherence of *K. pneumoniae* to host cell and establishment in mice, including liver, spleen, kidney and lung tissues, and induce the formation of liver abscess in mice. Our results also demonstrated that *K. pneumoniae bolA* increases the production of siderophore and virulence in Galleria mellonella larvae. Collectively, our results demonstrated that *K. pneumoniae* BolA is a new virulence factor which contributes to survival in different stresses and overcome host defense. These findings are helpful for the research of new treatment strategies for *K. pneumoniae* infection.

**IMPORTANCE:** *Klebsiella pneumoniae* is an important conditional pathogen causing nosocomial infections and community-acquired infections. It can resistant to multiple antibiotics, causing refractory infections and public health threat. Therefore, new treatments are required to fight the pathogen, and a better understanding of its virulence factors are needed to develop new drugs. Here, we unraveled the role of BolA in survival under different stresses and overcome host defense. Our results suggested that *bolA* actively contributes to cell morphology, stresses challenge, cell adhesion and siderophore production that are tightly related to bacterial virulence. Therefore, *bolA* mutant strain reduces the virulence of *K. pneumoniae* in G. mellonella larvae and its colonization ability in mice. These results reported *bolA* is a key virulence factor in *K. pneumoniae*, and they are helpful for research of new therapies to treat this increasingly problematic pathogen.

---

*Klebsiella pneumoniae* is an opportunistic pathogen that causes community infections and hospital infections^[1-3]^. It has the ability to metastasize spread, as well as extremely high morbidity and mortality^[4, 5]^. *K. pneumoniae* is classified into two types: classical *Klebsiella pneumoniae* (CKP) mainly occurs in hospitals and long-term-care facilities, while hypervirulent *Klebsiella pneumoniae* (hvKP) causes community-acquired, tissue-invasive *K. pneumoniae* infections in otherwise healthy individuals that often presented in a variety of sites^[6]^. These cases included severe pneumonia, urinary tract infection (UTI), sepsis, pyogenic liver abscess (PLA), endophthalmitis, meningitis and necrotizing fasciitis^[4, 7-9]^. *K. pneumoniae* ubiquitous in the environment and is often found in the gastrointestinal tract and on medical devices^[10, 11]^. The loss of porins, overexpression of efflux pumps, the prevalence of carbapenemases and extended-spectrum β-lactamases often lead to the emergence of multidrug-resistant *K. pneumoniae*, causing a variety of refractory infections ^[6][12-14]^. Therefore, in order to combat these daunting infections, it is necessary to identification of critical virulence factors in *K. pneumoniae*.

Environmental stresses can induce adaptive responses of bacteria cells to ensure survival, which is often related to virulence^[15, 16]^. Stress response protein BolA was first discovered in *E. coli*, and its homologues form a broadly conserved family of proteins in prokaryotes and eukaryotes^[17]^. In *E. coli*, it was discovered that *bolA* can make cells shorter and thicker after a period of starvation or stationary phase conditions, and the ability of bacteria to perception and adapt to harsh environmental conditions is critical to survival^[18]^. Later, it was found that the expression of *bolA* was affected by growth-rate, *bolA* induced expression during the transition from exponential phase to stationary phase due to the depletion of nutrients^[19]^. At the exponential phase, *bolA* expression is quickly induced in response to several stresses, such as sudden carbon starvation, osmotic stress, acidic stress, heat stress or oxidative stress, allowing the bacteria to quickly adapt to harsh environmental conditions^[20]^. In addition, *bolA* has an essential role in the regulation of cell permeability^[21]^ and biofilm formation^[22, 23]^. In recent years, further studies have found that BolA is a bacterial transcription factor, that elevated levels of BolA can induce tricarboxylic acid (TCA) cycle genes, fimbria-like adhesins-related genes and repression of flagellum-associated genes with consequences for bacterial motility and biofilm formation^[24]^. It also participates in the expression of DGCs and PDEs, which affects the synthesis and degradation of secondary signal metabolite c-di-GMP^[25]^. These studies prompted us to research the effect of transcription factor BolA on the virulence of *K. pneumoniae*.

*K. pneumoniae* have multiple virulence factors including lipopolysaccharide, capsule polysaccharide, outer membrane proteins, fimbriae, nitrogen source utilization and siderophores, for survival and immune evasion during infection^[9]^. It is worth noting that *K. pneumoniae* iron uptake genes are associated with capsule formation, Hypermucoviscosity (HMV), purulent tissue infection and are found in invasive strains^[26]^. In the present work, we identified a *bolA* homolog in *K. pneumoniae* NTUH-K2044 and unravel its role in several phenotypes associated with the process of cell morphology, siderophore production, cell adhesion, tissue colonization, liver abscesses and virulence.

## RESULTS

### BolA is required for *K. pneumoniae* cell morphology and siderophore production

To search for homologs of *K. pneumoniae* BolA, the *E. coli* K-12 W3110 amino acid sequence (GenBank accession no. APC50728.1), *S. Typhimurium* SL1344 amino acid sequence (GenBank accession no. FQ312003.1) and *K. pneumoniae* NTUH-K2044 amino acid sequence (Laboratory sequencing results) were used. The amino acid sequences of *bolA* in *K. pneumoniae* and *E. coli* are 91.4% similarity, and that of S. Typhimurium SL1344 was 91.4% (data not shown). Moreover, the predicted 3D structures of BolA proteins are nearly identical for the three species, (Fig. 1A). *K. pneumoniae* and *E. coli* BolA protein have a α1β1β2α2α3β3α4 fold with four α-helices and three β-sheets (Fig. 1B). We have used a virulent *K. pneumoniae* strain (NTUH-K2044) to construct a *bolA* mutant strain and complementation strain. Scanning electron microscopy of these cells revealed some effects of *bolA* on cell morphology (Fig. 1C). During exponential phase in LB medium, wild type (WT) cells exhibited the classical rod-shaped morphology, and during stationary phase in LB medium, WT cells exhibited shorter and wider morphology, but the Δ*bolA* cells were generally longer and thinner than the WT cells, and similar to WT cells shape during exponential phase. Moreover, the results of transcriptome sequencing showed that the “transport” genes were significantly enriched in GO enrichment analysis. And siderophores are an important virulence factor in *K. pneumonia*^[27]^. Therefore, we were interested in using CAS plate to detect the siderophores of *K. pneumoniae*. The WT strain exhibited a larger orange halo than bolA mutant strain (Fig. 1D). The *bolA* deletion reduced the area of the orange halo about 2.6 times that of the WT strain (Fig. 1E). This result suggests that the delete of the *bolA* gene reduces the production of siderophore. Nest, we investigated whether *K. pneumoniae bolA* affects biofilm formation. Using a 96-well plate model, our results showed that the biofilm biomass of *bolA* mutant strain was significantly lower than WT strain (Fig. 1F), suggesting that *bolA* is required for *K. pneumoniae* biofilm formation.

**FIG 1.**
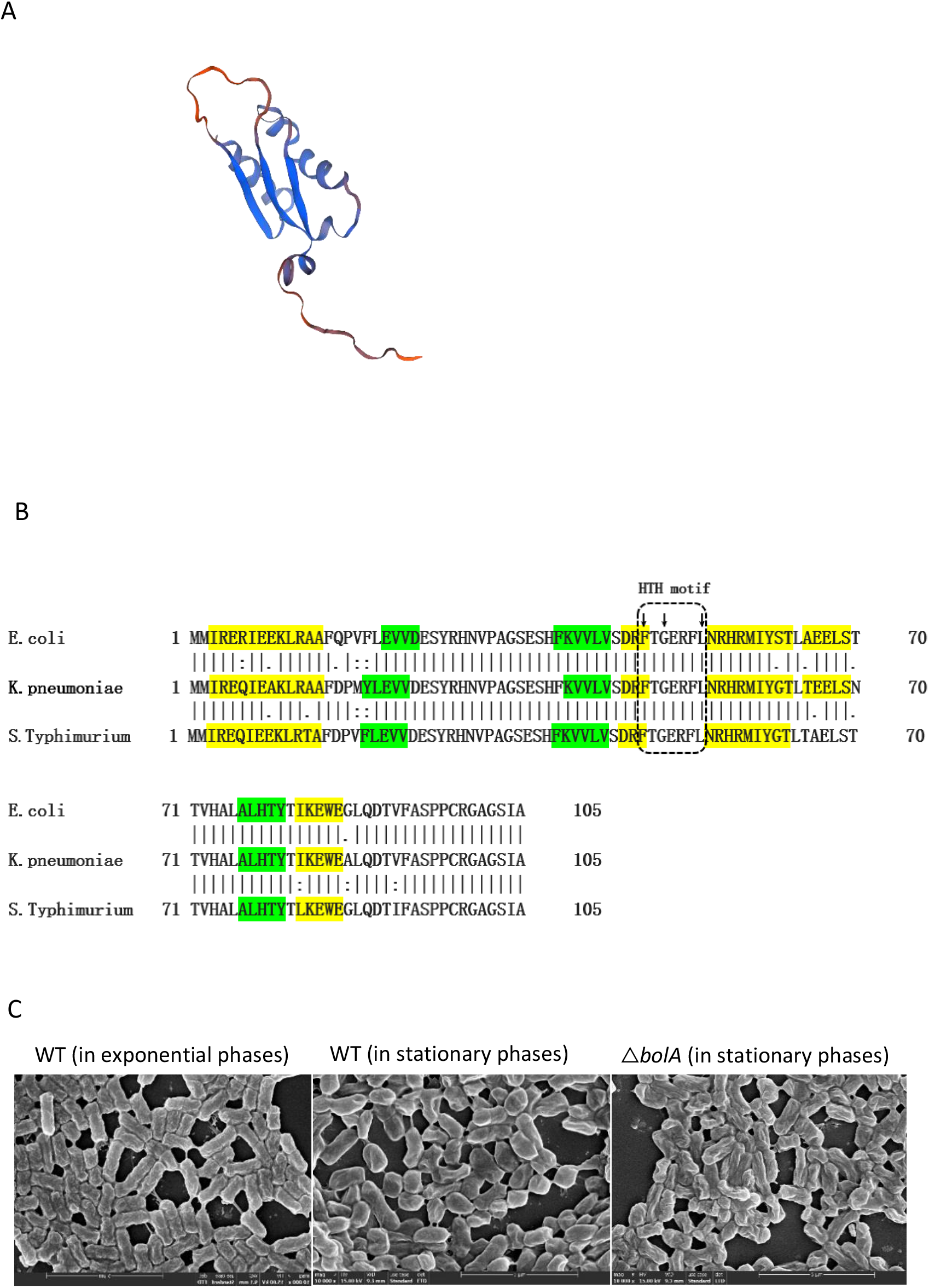
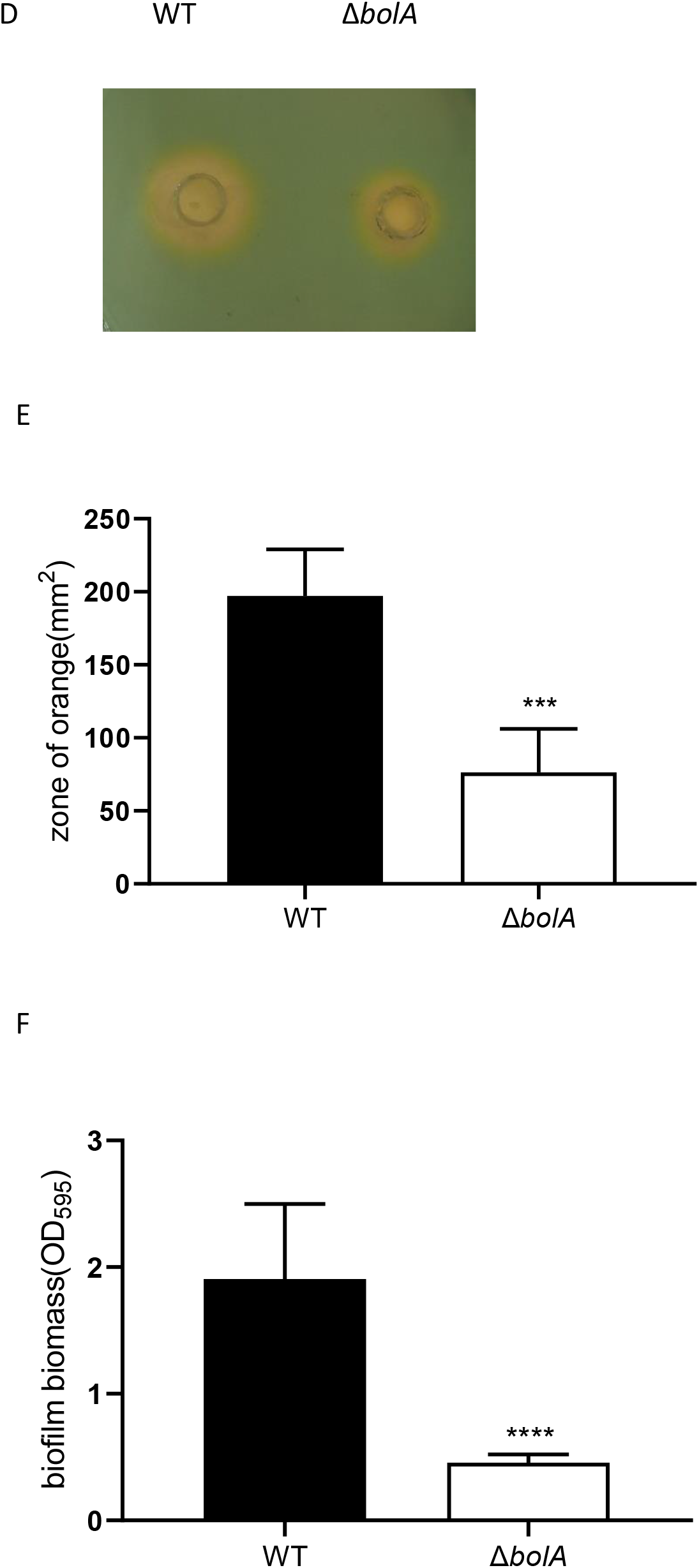
BolA is essential to the cell morphology of *K. pneumoniae*. (A) A computer model of the three-dimensional (3D) structure predicted from *K. pneumoniae* NTUH-K2044 BolA protein using SWISS-MODEL is presented. (B) Multiple sequence alignment of BolA from *E. coli* K-12 W3110, *S. Typhimurium* SL1344 and *K. pneumoniae* NTUH-K2044 obtained with the DNAMAN software. The secondary structure of three BolA proteins are predicted using SWISS-MODEL: in green are the β-sheets and in yellow are the α-helixes. The small arrows at the top of the sequence represent the amino acid of the characteristic helix to helix (HTH) FXGXXXL sequence. (C) Scanning electron microscopy images of the WT and *bolA* mutant strains bacterial cells are displayed. (D) The CAS ager plate was photographed after 48 h of cultivation at 28°C. (E) CAS plates were used for measure production of siderophore. The larger area of the orange halo indicates that the strain has produced more siderophores and increased the capacity of iron uptake. (F) The 96-well plate assay results showed *K. pneumonia bolA* mutant strain is defective in biofilm formation. After 48 h of incubation, the biofilm biomass was measured using 0.1% of crystal violet assay, and the optical density was determined at 595nm. ***, P < 0.001 by t test. ****, P<0.0001 by t test.

### BolA is an important bacterial transcription factor in *K. pneumoniae*

To evaluate the impact of *K. pneumoniae* BolA in global transcriptional regulation. Transcriptome sequencing was performed to compare the transcriptomic profiles of the WT and Δ*bolA* strains; 146 differentially expressed genes were revealed between these two strains in stationary phase. Of these differentially expressed genes, 26 were upregulated and 120 were downregulated. Most differentially expressed genes were distributed in downregulated.

Software GOseq based on Wallonia’s non-central hyper-geometric distribution were performed on the groups of genes significantly upregulated or downregulated. Among them, the down-regulated differentially expressed genes are mainly enriched in T2SS and T6SS, these two secretion systems are important virulence factors for bacteria. Gene Ontology (GO) enrichment analysis showed differentially expressed genes involved in membrane, localization, transport, oxidoreductase, tRNA processing (Fig. 2B). Kyoto Encyclopedia of Genes and Genomes (KEGG) enrichment analysis indicated differentially expressed genes involved in metabolism, transporters and secretion system (Fig. 2C). Stress response protein BolA was confirmed to be induced during the transition into stationary phase^[19, 28]^. It is interesting to observe that the genes in *bolA* mutant strain are downregulated during stationary phase, including several genes associated with secretion system *gspK*、*gspF*、gspE、*clpV*(*TssH*) and *VgrG*. More surprisingly, several genes associated with transport including *livF* 、 *livH* 、 *ydeY* 、 *KP1_3194* 、 *FbpB (KP1_3174)*、*fecD*、*FhuB (KP1_1440)*、*potC*、*potB*、*afuB*、*afuA*、*kdpB* and *kdpA* are downregulated (table 1) during stationary phase, in which *FhuB*、*fecD*、*FbpB*、*KP1*_*3194*、*afuB* and *afuA* genes are directly related to the iron transport. BolA also promotes the expression of several biofilm-related genes (*mhpB, ydeY, iolD*) and oxidoreduction-related genes (*ulaA*, KP1_4425, KP1_1420, KP1_2565).

**FIG 2.**
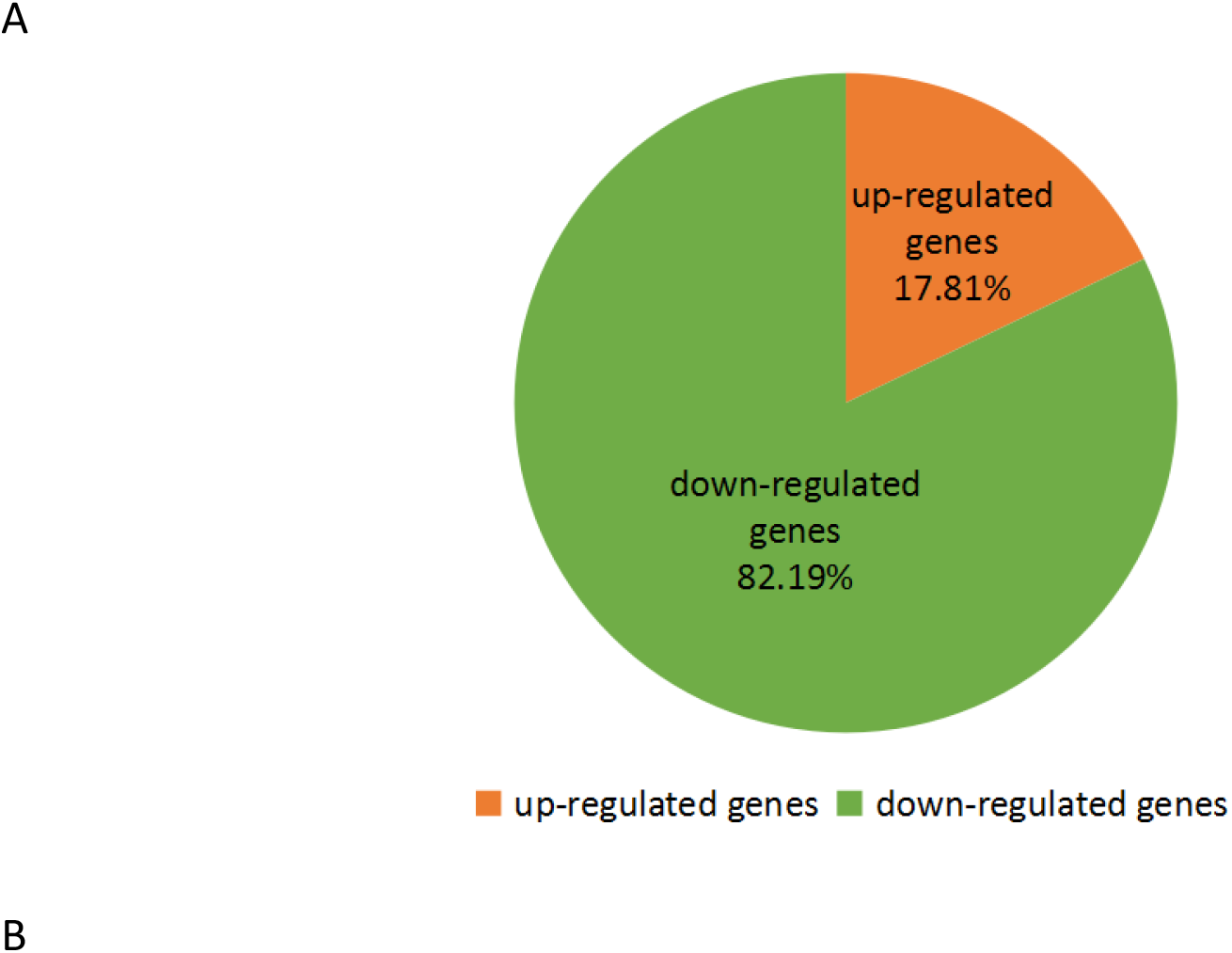
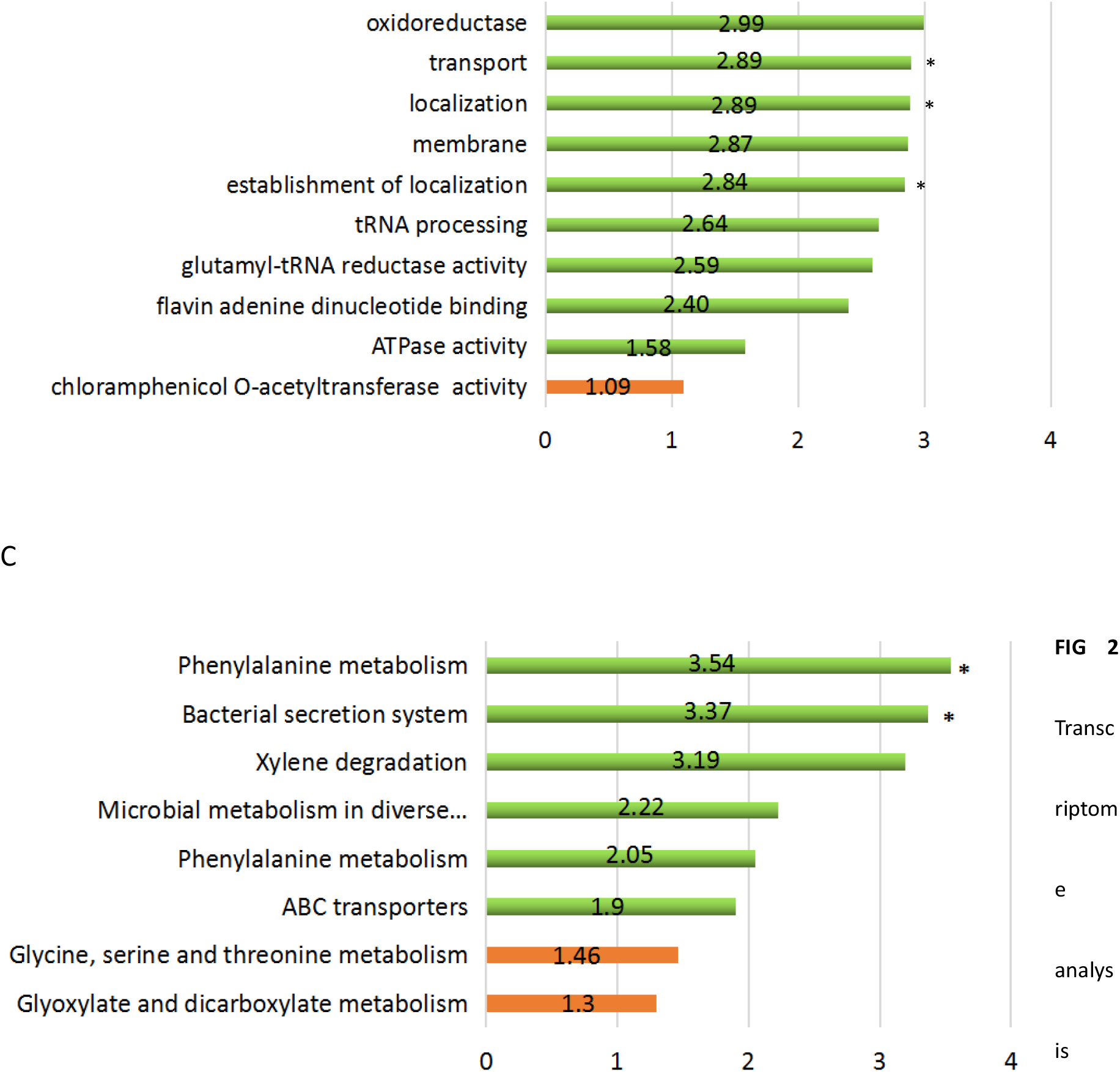
between WT and Δ*bolA* strains. (A) Pie chart of 146 genes significantly regulated by *bolA* gene. The number of genes significantly downregulated or upregulated are indicated in green and orange, respectively. (B) Graphical representation of the gene GO enrichment analysis is in “biological process,’’ “cellular component,” and “molecular function” categories for genes downregulated or upregulated in response to the *bolA* gene. *, P value associated with the enrichment test was lower than 1 × 10^−4^; No asterisk indicates that the P value associated with the enrichment test was between 1 × 10^−2^ and 1 × 10^−4^. (C) Functional classification of KEGG pathway. The KEGG enrichment pathway were summarized in eight main pathways. *, P value associated with the enrichment test was lower than 0.01; No asterisk indicates that the P value associated with the enrichment test was between 0.01 and 0.05. In all, the bar length corresponds to the average fold change (in absolute value of the WT /Δ*bolA* ratio) of the genes significantly upregulated or downregulated associated with each KEGG or GO enrichment pathways, and red graphic shows upregulated pathways and green the downregulated pathways.

**Table 1.** Differentially expressed genes involved in different pathways.

| Gene | log2.Fold_change. | P value |
| --- | --- | --- |
| Secretion system genes |  |  |
| *pulK* | -2.9395 | 0.00683 |
| *pulF* | -2.6985 | 0.00020409 |
| *pulH* | -4.4938 | 0.0026801 |
| *pulE* | -5.4198 | 1.76E-05 |
| *clpV* | -1.6083 | 0.001354 |
| *VgrG* | -3.0464 | 0.0017717 |
| Biofilm genes |  |  |
| *mhpB* | -4.4938 | 0.0026801 |
| *ydeY* | -2.3545 | 0.00017415 |
| *iolD* | -1.0834 | 0.00010384 |
| Oxidoreductase genes |  |  |
| *ulaA* | -4.3418 | 0.0046157 |
| *KP1_4425* | -3.4869 | 0.00011244 |
| *KP1_1420* | -2.4485 | 3.50E-06 |
| *KP1_2565* | -1.824 | 0.0021176 |
| K transport genes |  |  |
| *kdpA* | -1.824 | 0.0021176 |
| *kdpB* | -4.1932 | 8.04E-08 |
| Iron transport genes |  |  |
| *FhuB* | -2.3265 | 0.0027437 |
| *fecD* | -1.0755 | 0.0022485 |
| *FbpB* | -1.3265 | 2.57E-05 |
| *afuA* | -2.3265 | 0.0027437 |
| *afuB* | -3.9395 | 1.70E-06 |
| *KP1_3194* | -1.1371 | 3.85E-05 |

### BolA is an important metabolic modulator of *K. pneumoniae*

To compare the differentially accumulated metabolites between WT and Δ*bolA* strains, LC-MS was used for metabolite identification. In total, 82 differentially accumulated metabolites with known specific substances were screened. These differentially accumulated metabolites were defined as those with a variable importance for projection (VIP) value >1, P value <0.05 compared to WT. Among the detected metabolites, 25 metabolites were upregulated, and 57 metabolites were downregulated. Among them, 17 metabolites were increased, and 24 metabolites were decreased in pos mode, while 8 metabolites were increased, and 33 metabolites were decreased in neg mode. The differentially accumulated metabolites were visualized by volcano map (Fig. 3A).

**FIG 3.**
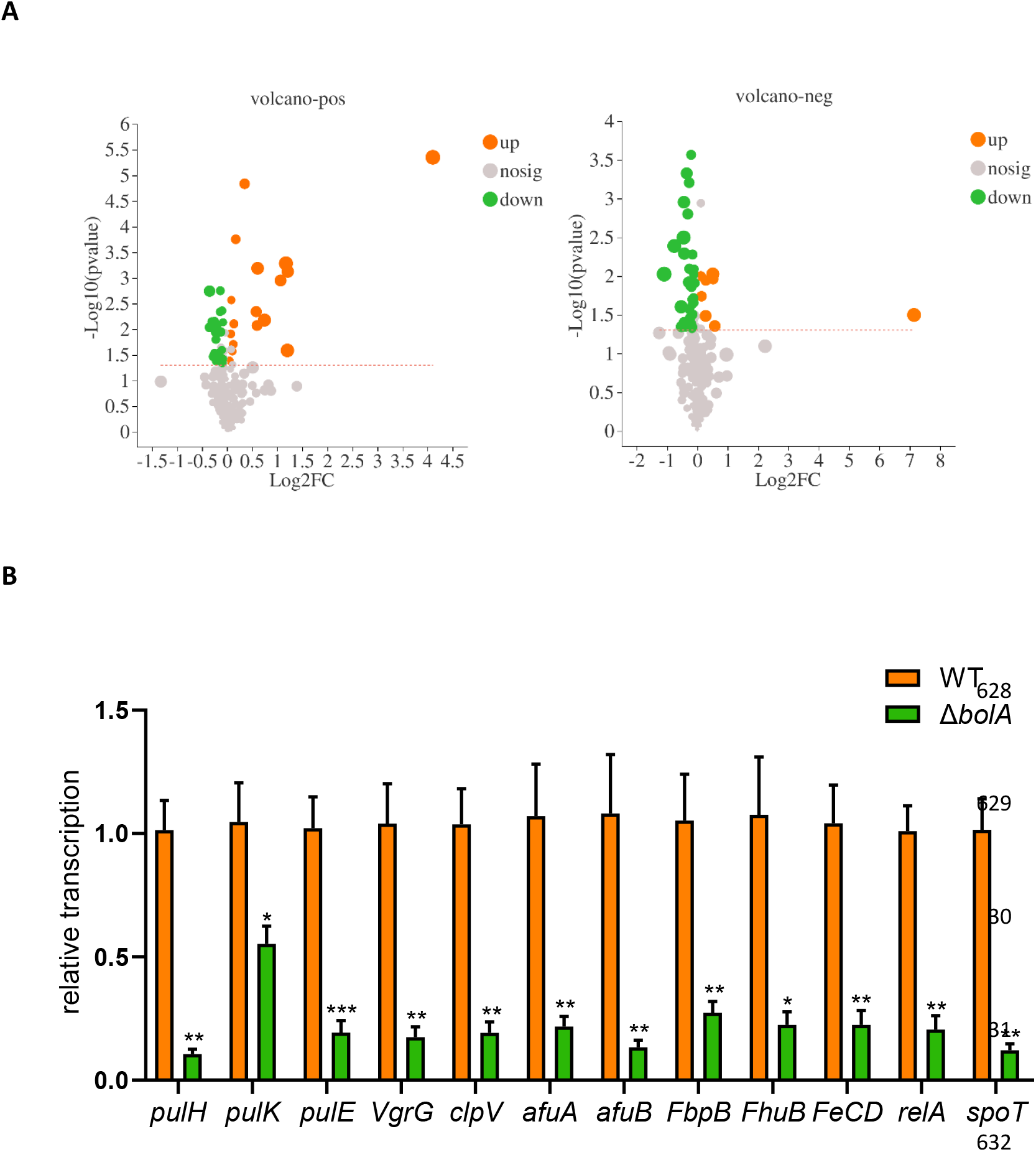
A. Volcanic maps of differentially accumulated metabolites in WT and Δ*bolA* strains. Red dots are upregulated metabolites, and green dots are downregulated metabolites. B. Results of qPCR tests are showed.

After metabolomic analysis, five metabolites related to stress resistance and virulence were identified, which were Agmatine, Cadaverine, Guanosine, Flavin adenine dinucleotide (FAD) and D-biotin respectively (table 2). It is speculated that the downregulation may be the factor leading to the weakening of the virulence and stress resistance of the *bolA* mutant strain of *K. pneumoniae*.

**Table 2.** Differentially accumulated metabolites of virulence and stress resistance.

| Name | Regulated type | FC(*bolA*/WT) | VIP | P value |
| --- | --- | --- | --- | --- |
| Agmatine | down | 0.87 | 1.28 | 0.0368 |
| Cadaverine | down | 0.91 | 1.17 | 0.0045 |
| Guanosine | down | 0.94 | 1.16 | 0.0454 |
| FAD | down | 0.80 | 1.84 | 0.0016 |
| D-Biotin | down | 0.79 | 1.53 | 0.0092 |

In order to validate these results, the expression of several of these genes were further analyzed by qPCR (Fig. 3B). Twelve different genes were selected for analysis, including the differentially expressed genes mentioned above and ppGpp synthetic-related genes. The regulation pattern of these genes is consistent with the transcriptome analysis results, and these genes are downregulated after *bolA* deletion.

### BolA promotes *K. pneumoniae* survival in bile and oxidative stresses

To evaluate the role of *K. pneumoniae bolA* in intestinal colonization, bacteria underwent specific intestinal stresses associated with osmotic, bile and oxidative. For the osmotic resistance assay, no significant difference was observed between the *K. pneumoniae* NTUH-K2044 and *bolA* mutant strains in LB medium containing 0.5M of NaCl (Fig. 4A). In bile challenge experiment, we were examined the ability of the strains to tolerate bile in LB medium, and significant difference was observed between the WT and Δ*bolA* strains (Fig. 4B). WT, Δ*bolA* and complementation strains were growth in LB medium containing 1% of ox bile, the ability of WT strain to grow in the 1% bile was three-fold higher than Δ*bolA* strain, and complementation strain restored the ability to tolerate bile stress. Oxidative stress assay showed that the Δ*bolA* strain had 1.79-fold greater sensitivity to 30% H_2_O_2_ (inhibition zone =13.73 cm^2^) than the WT strain (inhibition zone =7.69cm^2^) (Fig. 4C). Collectively, these results suggest that the response of the *bolA* in *K. pneumoniae* in bile, and oxidative stresses.

**FIG 4.**
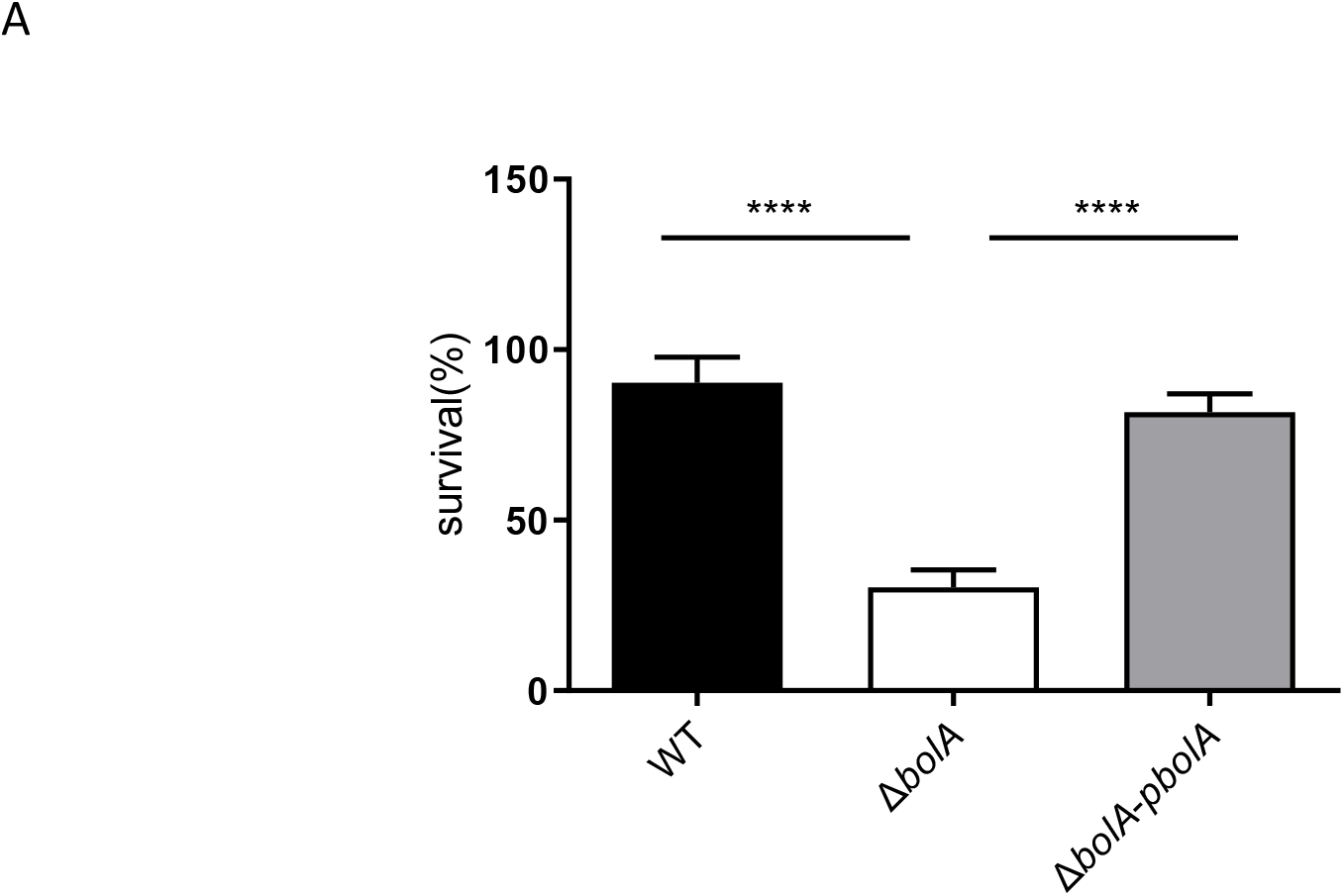
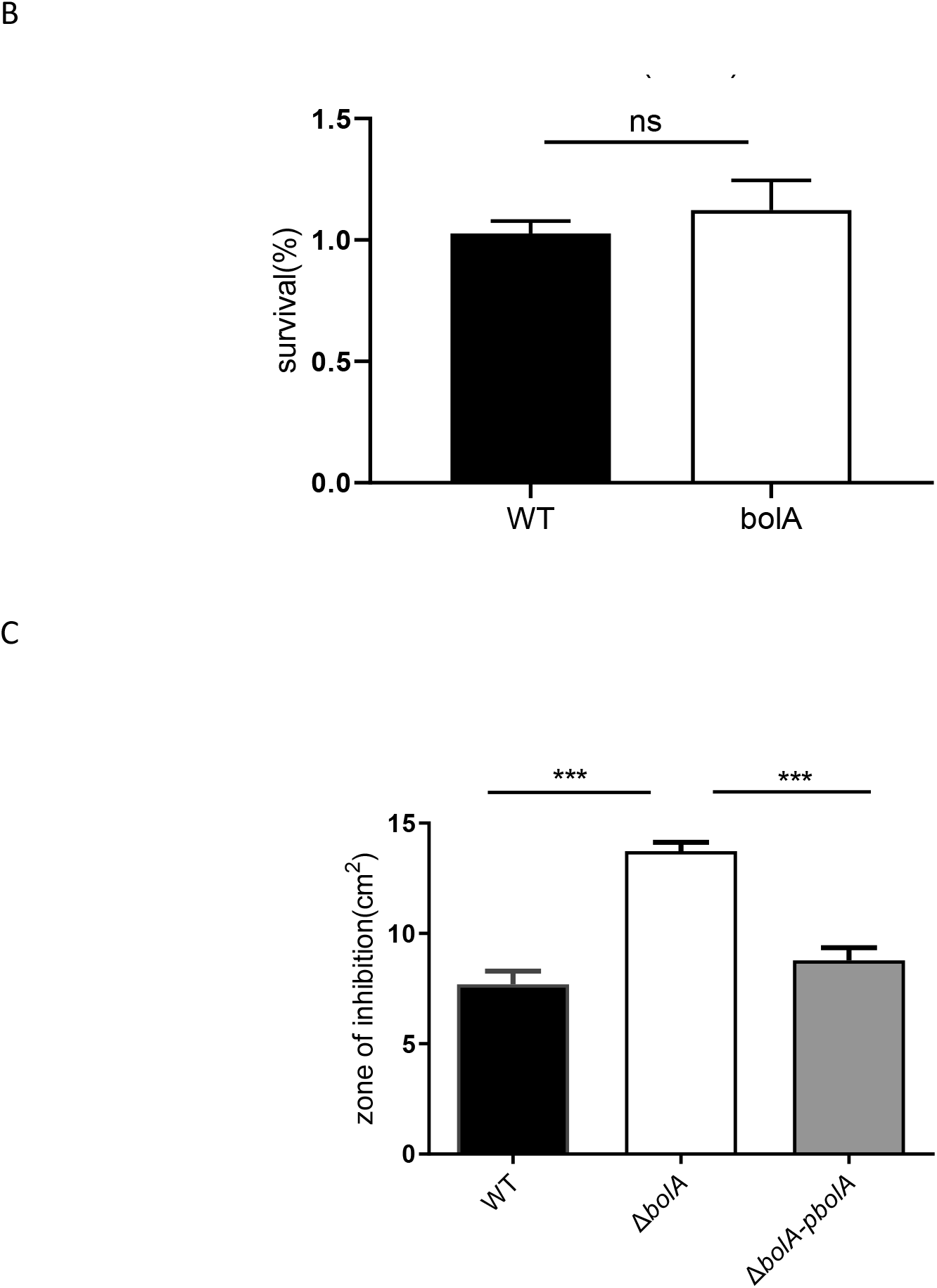
BolA augments *K. pneumoniae* resistance to osmotic, bile and oxidative stresses. (A) The percentage of viable cells were calculated by plating the appropriate dilutions on LB ager plates containing bile (1.0%). (B) The ability of resistance to osmotic stress for WT, andΔ*bolA* strains were extrapolated by comparison to the numbers of viable cells on LB ager plates containing NaCl (0.5 M). (C) The ability of strains to challenge hydrogen peroxide (30%) was measured by disc diffusion assay. ***, Significant difference (P<0.001, t test), ****, Significant difference (P<0.0001, t test), NS, not significant (P>0.05, t test

### BolA promotes *K. pneumoniae* cell adhesion and colonization of mice

The WT and Δ*bolA* strains were compared for their ability to adhere to HCT116 human colon cancer cells (Fig. 6). The *bolA* mutant strain was significantly decreased adherence compared with the WT strain, mounting 2.8fold lesser adherence to HCT116 cells. Next, we performed in vivo verification. We used a mouse septicemia infection model to observe the colonization of bacteria in mouse tissues Including the liver, spleen, lung and kidney. Ten-week-old female BALB/c mice were intraperitoneally injected with 2× 10^4^ CFU in 500μL of NS. The mice were euthanized at 24h postinoculation (hpi) and the liver, spleen, lung and kidney bacterial burdens were determined. Given transcription factor *bolA* deletion, we anticipated that the mutant strain would be attenuated, and this was indeed observed. We found that the colonization ability of *K. pneumoniae* in liver, spleen, lung and kidney tissues of mice after the *bolA* deletion was significantly lower than the WT strain. In the liver, spleen, lung and kidney at 24 hpi, the Δ*bolA* strain had colonization levels about 4 logs lower than *K. pneumoniae* NTUH-K2044 (Fig. 6B). These data indicated that the *bolA* mutant strain ability to survive and spread was attenuated in the mice.

**FIG 5.**
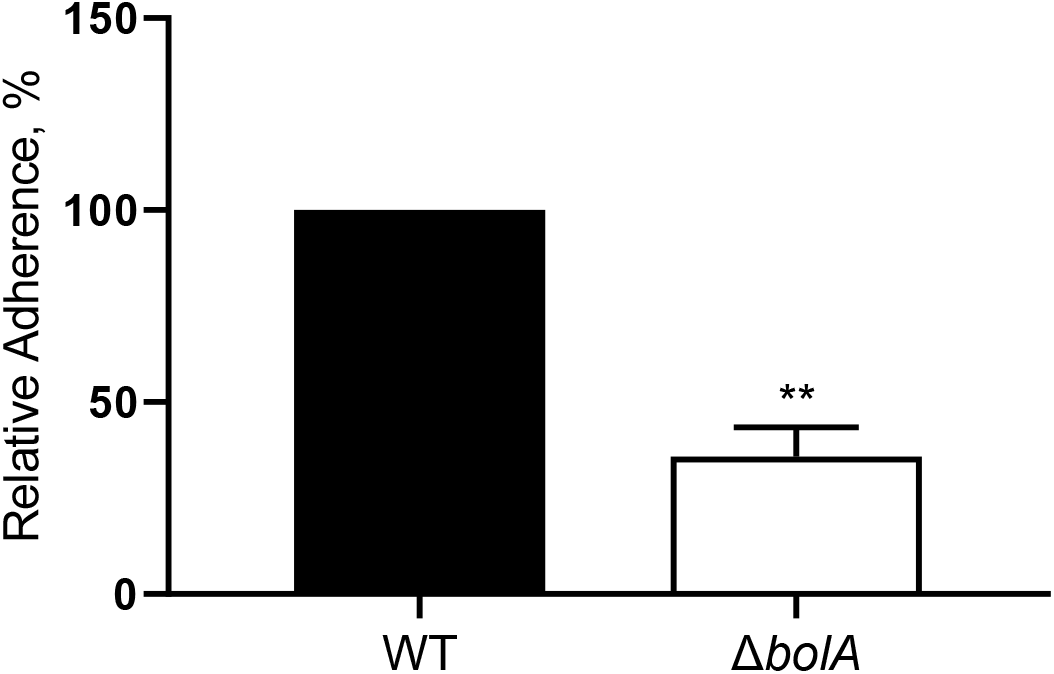
Adherence of *K. pneumoniae* NTUH-K2044 WT and Δ*bolA* strains to HCT116 cell. The logarithmic phase *K. pneumoniae* were added to cells of the HCT116. Next, bacteria were settled onto host cells by centrifugation at 200 × g for 5 minutes, and the mixture was incubated for 1 hour, washed with PBS and then the bacteria are released with Triton X-100, and plated onto LB agar for CFU counts. Rates of adherence of the *bolA* mutant strain was normalized to that of WT strain (100%). **, P < 0.01 by t test.

**FIG 6.**
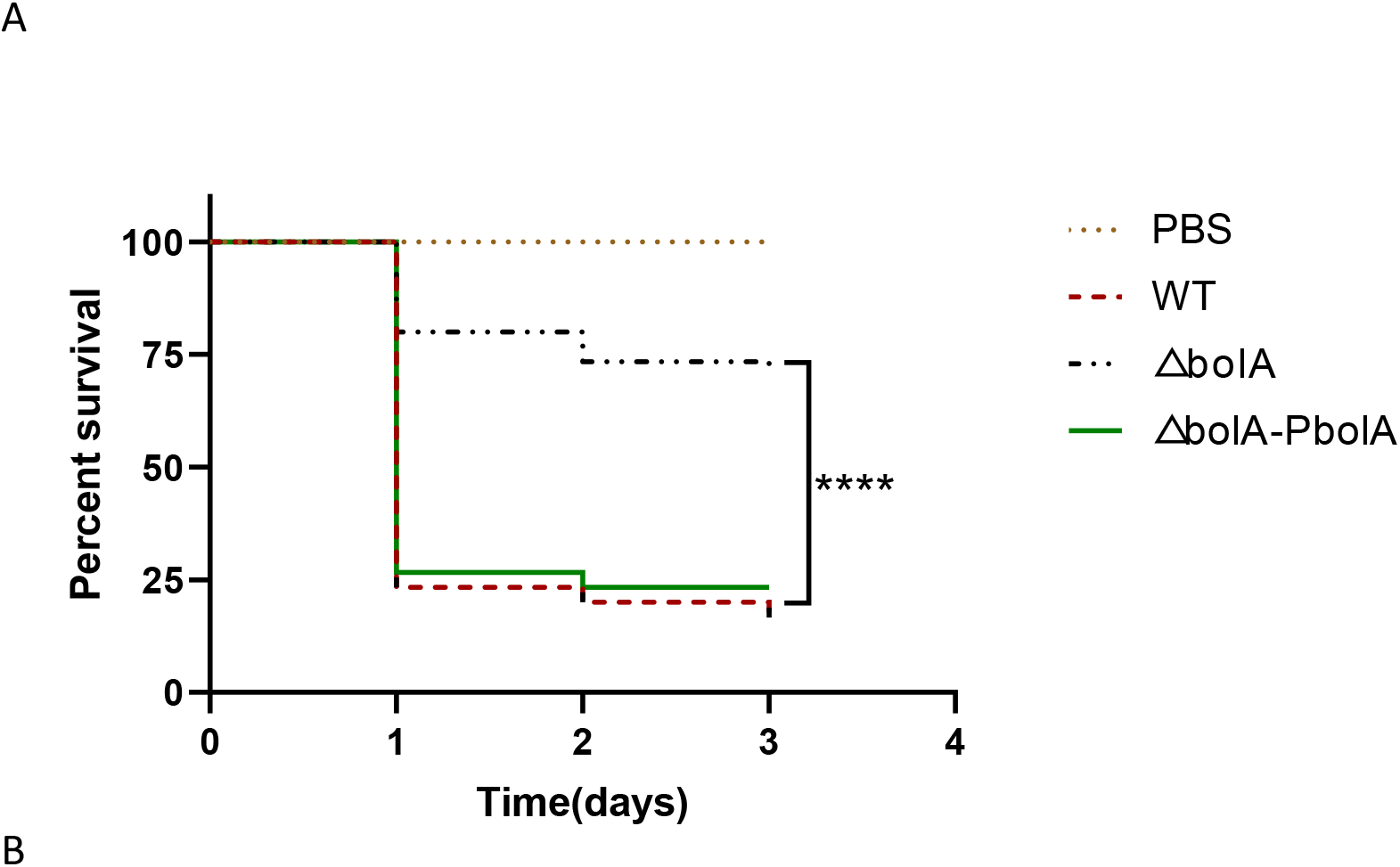
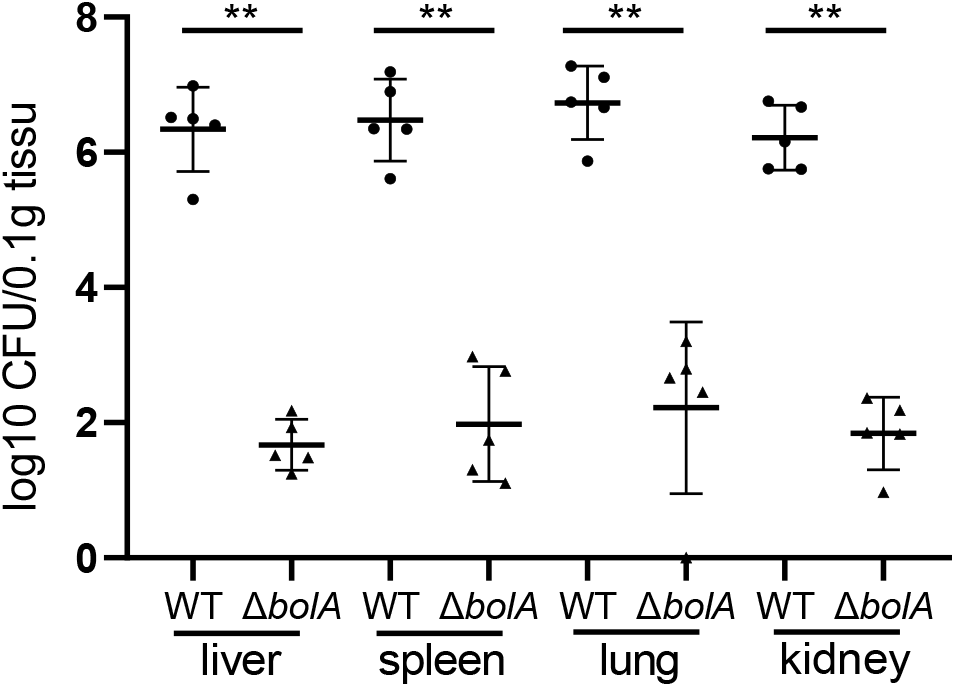
(A) *K. pneumoniae bolA* displays increased virulence in the G. mellonella larva infection model. Survival rate of larva infected with PBS or 1×10^7^ CFU of WT, Δ*bolA* and complementation strains. Larvae infected with Δ*bolA* strain had enhanced survival rate compared to *K. pneumoniae* NTUH-K2044 or the complementation strains. Statistical significance is examined by means of Kaplan-Meier analysis with a log-rank (Mantel-Cox) test (****, P<0.0001). (B) WT and Δ*bolA* strains were Intraperitoneal injection to ten-week-old female BALB/c mice (5 mice for each group) at an inoculation dose of either 1×10^4^ CFUs for 24 hours. The bacterial burden in liver, spleen, lung and kidney were determined for each mouse by quantitative plate count at 24 hours after infection. Bacterial numbers (expressed as log10 CFUs) were standardized per 0.1 gram of wet organ weight. Statistical significance is examined based on unpaired nonparametric Mann–Whitney U tests (**, P < 0.01).

Histopathological evaluation of the livers from WT-infected mice at 120 hours after infection revealed the frequent presence of either macroabscesses (Fig. 7C, arrow) or large abscesses (Fig. 7F, arrow). The high-power field (magnification, ×40) shows the microabscesses composed of inflammatory cells (Fig. 7D, arrow). In contrast, no liver abscesses were observed in Δ*bolA*-infected mice (Fig. 7E). The numbers of liver abscesses were 14, 17, 12, 3 (mean, 11.5) in 4 WT-infected mice, and 0, 0, 0, 0 (mean, 0) in 4 Δ*bolA*-infected mice (WT vs Δ*bolA*, P<0.01; Student t test). Moreover, In WT-infected mice, gross liver abscesses were observed (Fig. 7A, arrow). In conclusion, *K. pneumoniae* BolA can obviously induce the formation of multiple liver abscesses in mice.

**FIG 7.**
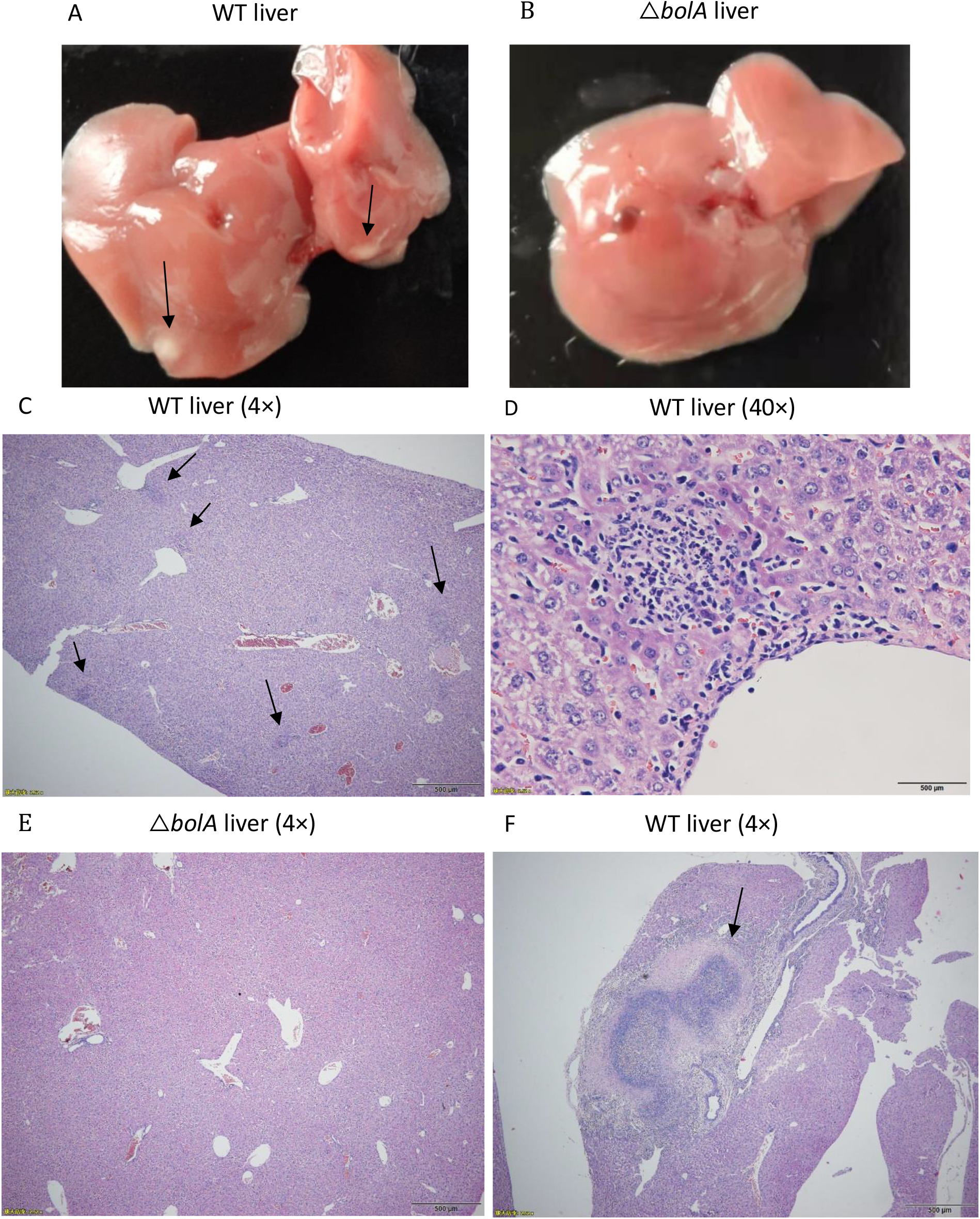
Histopathological examination of tissues from mice. Mice were inoculated with *K. pneumoniae* WT or *bolA* mutant strains (2×10^3^CFU). At 120 hours after infection, liver tissues were taken for hematoxylin-eosin staining sections, and representative microscopic photos were selected for display (Fig. 7). In mice infected with the WT strain, large abscesses (F, arrow) or microabscesses (C, arrow) were seen in the liver (magnification, ×4); The high-power field (magnification, x40) shows the microabscesses composed of inflammatory cells (D). Furthermore, in mice infected with WT strain, the resected liver show significant abscesses (A, arrow); No microabscess formation was observed in the liver (magnification, ×4) of mice infected with *bolA* mutant strain (E), and the resected liver show no obvious abscess (B).

### The BolA promotes *K. pneumoniae* virulence

To evaluate the involvement of *bolA* in virulence, we used the G. mellonella larvae for *K. pneumoniae* infection^[29]^. G. mellonella larvae were injected with PBS or 2×10^5^ CFU/larva of the *K. pneumoniae* NUTH-K2044 WT, Δ*bolA* or complementation strains, and the survival rate was daily monitored over a period of 72 hours. The results of infection by the WT strain were clearly seen at 72 hours of infection, with a decrease of about 83% of the initial larvae population (Fig. 6A). It’s worth noting that the *bolA* mutant strain was found to increase by 53% the larval survival rate at 72h of infection compared with the survival rate of the WT strain (Fig. 6A). Complementation of the *bolA* mutant strain was shown to majority restore the virulence to WT levels (Fig. 6A). These results clearly indicate the essential role of *bolA* in the pathogenesis of *K. pneumoniae* in the G. mellonella larvae infection model.

## DISCUSSION

In this paper, we characterized the *K. pneumoniae* NUTH-K2044 BolA protein. The BolA protein is highly conserved among *E. coli, K. pneumoniae* and *S. typhimurium*. In *E. coli*, when bacteria enter the stationary phase or under harsh environmental conditions, the *bolA* will be overexpressed to make the bacterial cells become rounder and shorter to protect the bacteria^[30]^. We saw the same results in *K. pneumoniae*. The cell morphology of the *bolA* mutant strain during stationary phase cannot be rounder and shorter like the WT strain. The shorter and rounder of bacterial cells result in a decrease in surface to volume ratio, and thus less surface area exposed to the damaging or unfavourable environments. This is consistent with previous research that overexpression of *bolA* induces formation of spherical cells in *E. coli*^[20]^. Growth analyses revealed that the deletion of *bolA* in *E. coli* was reported to increase the growth rate compared to the WT strain in a nutrient-rich medium (LB medium)^[17]^. However, the deletion of *bolA* in *S. Typhimurium* strain (SL1344) did not significantly affect the growth rate relative to the WT strain ^[28]^. In *K. pneumoniae* NTUH-K2044, while different, *bolA* mutant strain did not significantly impact the growth rate relative to the WT strain, and only a slight increase in the optical density at 595nm of the *bolA* mutant strain at the end of the exponent phase and the value of OD_595_ a slight decrease at the stationary phase was observed. However, in M9 medium, growth analyses of *bolA* mutant and WT strains revealed that the *bolA* deletion significantly reduced the growth rate and terminal density at 595nm (see Fig. S1). In addition, the researchers have found that the importance of accurate regulation of cross-talk between second messenger c-di-GMP and *bolA* for a proper response in the regulation of biofilm development^[25]^. Therefore, we also conducted biofilm formation experiments and MIC assays. While, *bolA* did not impact the MIC level (see Table S1). And the ability of biofilm formation of *bolA* mutant strain was significantly lower than WT strain, which is consistent with previous studies^[31]^. These results providing additional evidence that *bolA* plays distinct roles in these organisms.

BolA is a bacterial transcription factor that directly binds to the promoter region of a number of important genes, *flhD, fliA, gltA*, and *mreBCD*^[31]^. Several studies have indicated the critical contribution of transcriptional regulators in bacterial adaptation and virulence^[32-36]^. These studies emphasize the significance of the identification of new transcriptional regulators to understand and overcome *K. pneumoniae* pathogenesis strategies. In *E. coli*, BolA is a stress response protein whose expression increases under imposition of diverse severe conditions, conferring protection to the cells^[37]^. The deletion of *bolA* in *S. Typhimurium* was reported to increased survival under acid and oxidative stress compared to WT strain^[28]^. Furthermore, *bolA* expression is quickly induced in response to several stresses, such as sudden carbon starvation, osmotic stress, acidic stress, heat stress or oxidative stress, allowing the bacteria to quickly adapt to environmental changes at the log phase^[20]^. Above all, we suggested that BolA is a virulence factor that overexpresses itself in hostile environments to protect to the cells. These facts prompted us to research the effect of transcription factor BolA on the virulence capacity of *K. pneumoniae*. Unsurprisingly, our findings suggest that the response of *K. pneumoniae bolA* gene in bile, and oxidative stresses and increases its viability under both stresses. Our research also found that the deletion of *bolA* in *K. pneumoniae* significantly reduce its adhesion rate in intestinal cells HCT116. These studies point to a possible role of *bolA* in *K. pneumoniae* virulence capacity.

Interestingly, some results have showed that transcription factor BolA affects different pathways directly related to virulence^[17, 31]^. Iron is an essential nutrient for the host and most microorganisms^[38]^. Microorganisms produce siderophore which is a small molecule of iron chelators and is an important virulence factor and whose main function is to iron uptake from the host and provide this essential metal nutrient to microorganisms^[4, 39]^. Moreover, two characteristics known to distinguish CKP from hvKP strains are the number of siderophore iron acquisition systems and the abundance of capsules^[8]^. Classical strains usually have one or two siderophore iron acquisition systems, while HV strains have three or four, and HV strains produced very thick capsular and showed a high mucinous viscosity (HMV). In this study, transcriptome data and qPCR suggested BolA direct effects are related to the induction of genes related with the iron transport. So, we carried out CAS agar plate experiment and found that *bolA* has a positive regulatory effect on siderophore production. The results of research show that the *bolA* mutant strain exhibited litter halo than WT strain in a CAS ager plate. So far, this is the first time we have identified the role of *bolA* in capacity of Iron uptake. We hypothesized that *bolA* affects the production of siderophore by regulating iron transport-related genes.

In the field of infection biology, G. mellonella larvae are becoming an attractive infection model for human pathogens. G. mellonella larvae can grow at 37°C, so the model of larvae is more suitable for the study of human pathogenic bacteria. And it has innated immune like that of mammals’ response to infections. In G. mellonella larvae killing experiment, our results suggest that the delete of the *bolA* gene significantly reduced the virulence of *K. pneumoniae* in the larva of G. mellonella. The results showed that *K. pneumoniae bolA* gene was absent and the pathogenicity of the bacteria was decrease, similar to what was observed for the *S. Typhimurium bolA* homolog^[28]^.

In addition to *bolA* effect on cell morphology, growth rate in M9 medium, resistance to bile and oxidation also influenced its capacity to colonize in mice and induced the formation of liver abscess in mice. In *V. cholera, ibaG* is a *bolA* gene’s homologue. Very recently, we have shown that the deletion of *ibaG* in *V. cholera* exhibited obvious colonization defects in intestinal tract colonization. When stationary-phase cultures were used to infect mice, the *ibaG* mutant strain showed 50-fold deficit in colonization compared with WT strain inoculated^[40]^. Thus, we hypothesized that the *bolA* deletion significantly reduced the ability to colonize the liver, lung, spleen and kidney organs of mice. Not surprisingly, we observed the same results for the first time, showed that *bolA* induced the formation of multiple microabscesses and large abscesses in mice liver tissue. Its ability to adhere to the intestinal epithelium is attenuated. Adhesion to intestinal epithelial could happen in the earlier stage of infection, seem to be important for *K. pneumoniae* pathogenesis. Histopathological examination revealed that the deletion of *bolA* in *K. pneumoniae* exhibited a lack of ability to induce liver abscess. These data indicate that *bolA* is essential for *K. pneumoniae* to survive, colonize, which contributes to liver abscess generation and host death. We hypothesized these are related to that the *bolA* mutant strain is associated with decreased colonization ability in the host and reduced the ability of bacteria cells to protect themselves in the host.

Transcriptome results showed that the expression of T2SS, T6SS and iron absorption-related virulence genes of the Δ*bolA* strain were significantly down-regulated. In addition, the metabolome results showed that related to virulence and stress resistance of the Δ*bolA* strain metabolites (including biotin, spermine, cadaverine, guanosine, and FAD) were downregulated. And the results of qPCR tests showed that the expression of T2SS, T6SS, iron transport and ppGpp synthetic related genes were downregulated after *bolA* was deleted, the results of transcriptome sequencing were consistent. It is worth noting that guanosine can become c-di-GMP when phosphorylated. Furthermore, c-di-GMP and ppGpp are second messenger of bacteria that plays an important role in signal transduction. C-di-GMP can regulate a variety of biological pathways. Previous studies have found that the increase in the level of c-di-GMP has a significant inhibitory effect on the movement of *Pseudomonas aeruginosa*, positively regulates the formation of biofilms and the production of siderophore, and improve resistance to oxidative stress^[41]^. The ppGpp has a variety of physiological functions, the most important of which is to coordinate the stress response of bacteria to stress, and the RelA/SpoT homologous protein is a key protein that regulates the level of ppGpp^[42]^. Therefore, we hypothesized that virulence factor BolA may promotes the virulence of *K. pneumoniae* by promoting the accumulation of five metabolites (including biotin, spermine, cadaverine, guanosine, and FAD) and the expression of genes related to T2SS, T6SS, iron transport and ppGpp synthesis.

In order to investigate the different roles of *bolA* of *K. pneumoniae*, we tested the different strains of iron uptake, M9 medium and LB medium to growth rate, the adhesion of different strains to intestinal cells HCT116, their resistance to bile, osmotic and oxidative stress, as well as the ability of killing G. mellonella larvae. Finally, the colonization ability of different strains in liver, lung, spleen and kidney tissues of mice and the role of *bolA* in liver abscess formation were tested by mice septicemia infection model. Taken together, we demonstrate, for the first time, that *K. pneumoniae* virulence factor BolA has important regulatory roles in bacterial morphology, biofilm formation and production of siderophore and virulence. In particular, the *bolA* gene was deleted, which reduced bile and oxidative stress resistance and reduced liver, spleen, lung and kidney colonization in mice. This study show that virulence factor BolA is a good candidate as a therapeutic target against *K. pneumoniae* systemic infection.

## MATERIALS AND METHODS

### Bacterial strains, plasmids, primers, and growth conditions

The bacterial strains and plasmids used in the present work are listed (see Table S2). All the primers used in this study are listed in Table 2. All bacterial strains were stored at −80°C in lysogeny broth (LB) medium containing 25% glycerol. pkO3-Km was used to create gene mutant strain by homologous recombination. Unless otherwise stated, *K. pneumoniae* and *E. coli* cultures were grown in LB medium, M9 medium or on LB ager at 30°C, 37°C or 43°C and supplemented with Kanamycin (Km, 50μg/ml or 25μg/ml) where required. Bacterial growth was monitored by measuring the optical density at 595nm.

### Construction of the *bolA* gene deletion and complementation strains

A *bolA* gene deletion strain of *K. pneumoniae* NTUH-K2044 strain was constructed with the previously described method^[43]^. In brief, the left and right flanking DNA fragments of *bolA* gene were amplified by overlap PCR and cloned into pKO3-Km by digested, ligated, and then introduced into NTUH-K2044 by electroporation. To construct the complementation strain (Δ*bolA*+*bolA* strain), the DNA fragment that contained the *bolA* coding sequence and promoter region (using primers *bolA*-HB-F/R) (see Table S3) was amplified by PCR. Then, the DNA fragment was cloned into the pGEM-T-easy-km. After, the recombinant plasmid was transformed in Δ*bolA* strain by electroporation. The gene deletion mutant and complementation strains were verified by PCR and sequence determination.

### Scanning electron microscopy

The supernatants were discarded after bacterial cultures were centrifuged. Bacterial cells were overnight fixed at 4°C in 2.5% glutaraldehyde. Followed by washing them three times with PBS, and then dehydrated by gradient incubation, and finally anhydrous ethanol was used for suspension. Drops of the bacterial suspension were applied to a glass coverslip, dried, and covered with chromium. Finally, bacterial cells were observed and photographed.

### CAS agar assays for iron uptake

A chrome azurol S (CAS) agar assays were performed as previously described^[44, 45]^. In brief, 60 ml of CAS solution was first prepared by dissolving 60.5 mg CAS powder (Yuanye Bio-Technology, China) was added to 50 ml of double-distilled water (ddH2O), followed by 10 ml of 1 mM solution of FeCl_3_ (Aladdin) was added. Then, 72.9 mg hexadecyltrimethylammonium bromide (HDTMA; Yuanye Bio-Technology, China) was added in 40 ml ddH2O. At last, a 100 ml CAS stock solution was made by HDTMA solution was poured slowly with stirring into CAS solution and autoclaved to sterilize. Next, the freshly autoclaved 1.5% agar LB plate and CAS stock solution were mixed at a ratio of 9:1 to make A CAS agar plate. then, 2μl overnight bacterial culture (WT and Δ*bolA* strains) was inoculated to the CAS plate and cultivated at 28°C. The CAS plate was photographed after 48h of cultivation. The assay was repeated three times.

### Transcriptome sequencing

Overnight cultures were diluted 1:100 in fresh LB medium. Bacterial cultures were collected at stationary-phase. Then, the cells were harvested to extract total RNA. The total RNA was extracted, the quality of the RNA samples was detected, and Illumina sequencing was performed at Novogene Bioinformatics Technology Co., Ltd (Beijing, China).

### Assay Metabolome

Overnight cultures were centrifuged at 8000g for 10 minutes at 4°C. The supernatants were discarded; cell pellets were washed three times in 10 mL of PBS (4°C) and resuspended in an extraction solvent consisting of methanol/water (4:1 v/v), and a 6 mm grinding bead was added. After grinding for 6 minutes with a frozen tissue grinding instrument, the sample was extracted with low temperature ultrasound for 30 minutes, and then the sample was placed at −20°C for 30 minutes The supernatant was centrifuged for 15 minutes and transferred to a tube for analysis. The sample extracts were analyzed using an (Liquid chromatography mass spectrometry) LC-MS system, and the data were analyzed on the free online platform of Majorbio Cloud Platform.

### qPCR assays

WT and *bolA* mutant strains were grown in LB medium to stationary phase, then and bacterial cells were collected. The total RNA was extracted from harvested cells with Spin Column Bacteria Total RNA Purification Kit (Sangon Biotech). The cDNA was synthesized using TransScript® All-in-One First-Strand cDNA Synthesis SuperMix for qPCR (One-Step gDNA Removal). All qPCR assays were performed at least three times using Tip Green qPCR SuperMix (TransGen Biotech). The primers used for qPCR are shown (see Table S3). The data were analyzed by the ΔΔCT method using 16S mRNA as an internal control. Fold change was calculated using the ΔΔCT method. All qPCR reactions were performed in triplicate.

### Minimum Inhibitory Concentration (MIC) assays

MIC assays were performed by micro-dilution broth method in 96-well microplates. In short, bacterial cultures in exponential phase were diluted to a final OD_595_ of 0.1, then diluting it 1,000-fold into fresh LB medium. Next, 100μL of microbial suspension was added into each well and 100μL of antimicrobial agents were added only in the first well. From the first well, serial twofold dilutions of the antimicrobial agents were done and the plates were incubated without shaking for 24 h at 37°C. *E. coli* ATCC 25922 was used as a reference strain (control).

### Biofilm assays

The overnight cultures of bacteria were diluted to a final optical density OD_595_ of 0.1 in fresh LB medium. Then, 200μl of the diluted cultures were added to a 96-well Polystyrene (PS) plate and incubated at 37°C for 48 h. Bacterial cells were washed twice with ddH_2_O to remove plankton, and the attached bacteria were stained with 0.1% crystal violet for 10 minutes. Finally, the plate was washed four times with ddH2O. The crystal violet was solubilized by anhydrous ethanol, and the biofilm thickness was estimated by measuring the OD_595_ values.

### Stresses challenge assays

Bile, osmotic stresses assays were performed based on a previously described method^[46]^. Briefly, bacterial cultures were grown separately until they reached an OD_595_ of 0.2 in LB medium. Then, cultures were spread plated onto LB agar plates containing NaCl (0.25M, 0.5 M, 0.75M) and bile (1.0%, Sangon Biotech) respectively. The plates were incubated overnight at 37°C, and the numbers of colonies were counted. The results are expressed as the ratio of the number of colonies obtained from LB agar plate containing NaCl or bile to the number of colonies obtained from LB agar plate. These experiments were performed at least three times.

For oxidative stress sensitivity assays, the exponential-phase bacterial cultures were diluted to an OD_595_ of 0.2 and were uniformly spread over an LB agar plate. Then, a sterile 6-mm paper disk (5μl of 30% H_2_O_2_) was placed at the center on the agar surface. Next, The plates were incubated at 37°C for 24 h. The experiments were repeated at least three times.

### Adherence assays

HCT116 human colon cancer cells were grown in RPMI 1640 medium supplemented with 10% fetal bovine serum (FBS), 100 U/ml penicillin, and 0.1 mg/ml streptomycin^[47]^. Cells adhesion assays were performed mainly according to the methods described previously with minor modifications^[48, 49]^. For adherence experiments, HCT116 cells were grown to confluent monolayers in 24-well plates, and then the cells were washed twice with HBSS. Followed by 200μl of *K. pneumoniae* cells (OD_595_, 0.5) in FBS-free RPMI 1640 medium were added to each well and were settled onto host cells by centrifugation at 200×g for 5 minutes. After 1h of incubation in a humidified 5% CO_2_ mosphere at 37°C, cells were washed 4 times with PBS and then bacteria were released by the addition of 0.5% Triton X-100. Recovered bacteria were quantified by means of serial dilution and plating onto LB agar. The experiment was performed at thrice.

### G. mellonella larvae killing assays

G. mellonella larvae killing assays were performed as described previously^[50]^. In brief, *K. pneumoniae* strains were prepared by harvesting exponential phase bacterial cultures, washing twice in PBS and then adjusting to 1×10^7^ CFU. Groups of larvae (N = 10) were injected with 10μl of working bacterial suspension at the last proleg using a hypodermic microsyringe. For each assay, a group of larvae was injected with PBS as a control. Injected larvae were placed in petri dishes at 37°C for 72h in the dark. The percent survival was recorded at 24h intervals. G. mellonella larvae were considered dead if they did not respond to physical stimuli. Assays were repeated at three times.

### Mice infection experiments

Ten-week-old female BALB/c mice (n=5 in each group) were used for infections. Bacterial cells were collected by centrifugation and resuspended in sterile normal saline (NS). Mice were infected by intraperitoneal injection of the bacterial suspension (1× 10^4^ CFU *K. pneumoniae*/mouse). After 24h, the mice were euthanized and organs (including liver, spleen, kidney, lung) were homogenized in NS, and then appropriate dilutions were plated onto LB agar for CFU counts.

Eight-week-old female BALB/c mice (4 per group) were infected by intraperitoneal injection with *K. pneumoniae* at a dose of 2 × 10^3^ CFUs per mouse for 120 hours. The livers were retrieved, fixed in 10% formalin, and embedded in paraffin blocks. The tissue sections were stained with hematoxylin-eosin and were imaged and quantified under a light microscope. The number of abscesses was quantified in five low-power fields (magnification, ×4).

### Statistical analyses

All statistical analysis was performed using Prism (GraphPad). G. mellonella larvae survival was calculated by means of Kaplan-Meier analysis with a log-rank (Mantel-Cox) test. Differences in *K. pneumoniae* burden in tissues between groups were examined based on unpaired nonparametric Mann–Whitney U tests. Adherence assays, stresses challenge assays and iron uptake assays were analyzed by t test. Statistically significant was defined by P < 0.05 (*), P < 0.01 (**), P < 0.001 (***), and P < 0.0001 (****).

## ACKNOWLEDGMENTS

This research was funded by the National Natural Science Foundation of China [31500114] and the Sichuan Province Science and Technology project [2020YJ0338].

